# Time scales of mixing in an imperforate scleractinian coelenteron

**DOI:** 10.1101/2020.09.03.281709

**Authors:** Sara D. Williams, Mark R. Patterson

## Abstract

Coelentera are the largest components by volume in the gastrovascular system connecting polyps in a scleractinian colony. Thus to understand colony connectivity which is predicted to affect corals’ response to environmental change, we must first describe the dynamics inside these gastric cavities of individual polyps. We determined key time scales of mixing in coelentera by using microelectrodes to measure oxygen concentration after a light-to-dark transition in three polyps each of three colonies of *Montastraea cavernosa* in the laboratory. The gastrovascular system was modeled as an electrical network where voltage represents oxygen concentration, current represents oxygen flux, capacitors represent volume compartments, and resistors represent impedance to oxygen flux. The time constant of mixing, defined as the time needed for the system to disperse 63.2% of the fluid in the coelenteron, was determined from the oxygen dynamics in the coelenteron as modeled by a resistor-capacitor network. Time constants were on the order of three minutes and oxygen dynamics were well fit by the model prediction. However, as polyps depleted oxygen, we observed small magnitude (~ 0.1 ppm), high-frequency fluctuations in oxygen concentration. A power spectral density analysis identified two time scales of high-frequency mixing in the coelenteron. The greatest variance occurred at a period of 48.3 ± 2.8 seconds, with a secondary peak seen at 35.9 ± 2.3 seconds. The microenvironment within polyps of *M. cavernosa* can respond as fast or faster than their external environment can fluctuate, thus scleractinian polyps have the capacity to mediate their response to changing environmental conditions.

## Introduction

Internal fluid dynamics and mixing within the coral gastrovascular system are largely unknown but may be important for understanding coral response to environmental stressors. The level of connectivity among polyps, both structural via physical connections and functional via fluid transport dynamics, is understudied and likely to have important ramifications for all biochemical processes occurring in the colony. However, before we can understand the gastrovascular system at large, we first have to describe the dynamics within individual polyps as the coelentera are the largest components by volume. Ventilation periods and flushing rates of actinians (sea anemones), closely related to scleractinians, have been described (Jones et al., 1977; Nicol, 1959), but surprisingly the residence time of fluid and mixing rates in polyps have been largely ignored in investigations of scleractinian physiology.

Internal fluid movement has been well studied in hydroids and some octocorals (Blackstone, 1996; Harmata et al., 2013; Parrin et al., 2010; Gateno et al., 1998). *Acropora cervicornis* is one species of scleractinian coral with a well described perforate gastrovascular system (Gladfelter, 1983). This perforate coral has a flagella-lined gastrovascular system operating at a low Reynolds number, with viscous flow connecting polyps through a vast network of canals (Gladfelter, 1983). What little is known about mixing and the residence time of fluids within the main gastric cavity of scleractinians, the coelenteron, has been determined from studies of much larger, solitary anthozoan polyps. *Metridium senile*, a solitary anemone, controls the volume of fluid within its coelenteron through muscular expansion and contraction and intakes water through the inward-beating motion of cilia lining the siphonoglyph (Batham and Pantin, 1950). An individual *M. senile* can refill itself within a few hours after ‘shriveling’ (Batham and Pantin, 1950), and has a respiratory rhythm of periodic deflections caused by the release of coelenteric fluid through the mouth of 24-43 minutes (Jones et al., 1977). Metabolic activities that alter water chemistry inside a polyp’s gastrovascular system require periodic turnovers of the enclosed fluid, but this phenomenon has yet to be measured in corals.

An individual scleractinian polyp is essentially composed of a tubular, muscular pharynx leading to the coelenteron. These regions are subdivided by mesenteries, muscular folds of gastrodermis that are arranged “like spokes on a wheel in cross section” (Duerden, 1902) and function to increase surface area for digestion and metabolite exchange. Fluid is mixed in the coelenteron by muscular contraction, but mostly by ciliary action, as it has been estimated that every endodermal cell has one cilium (Gladfelter, 1983). Diffusion processes are limited at the coral surface, and in the gastric cavity, due to mucus production increasing the local viscosity (Patterson et al., 1991). Consequently, local fluid and mass transfer dynamics are driven by polyp behavior and active mixing.

Ventilation and mixing of the gastrovascular system affect oxygen and other metabolite dynamics in the coral colony. Oxygen is stratified within the gastric cavity, suggesting little to no exchange of fluid with the surrounding environment, creating a micro-environment inside the polyp (Agostini et al., 2012). This microenvironment has high levels of bacteria and nutrients and serves as a source of nitrogen and phosphate for photosynthetic activity (Agostini et al., 2012). The processes of host calcification and symbiont photosynthesis are tied through competition for available carbon (Jokiel, 2011), much of which takes place in the gastric cavity, surrounding endodermal tissue, and underlying calcifying fluid (Comeau et al., 2013; Cai et al., 2016; Bove et al., 2020). Host and symbiont metabolism strongly regulate the internal microenvironment of a coral polyp in addition to diffusional transport of materials from the environment. Carpenter and Patterson (2007) demonstrated intra-colony variation in metabolism caused by unidirectional flow over colonies of *Montastraea (Orbicella) annularis*, a species that also showed differential expression of heat shock proteins within a colony (Carpenter et al., 2010). Metabolic gradients in corals are assisted by the gastrovascular system and typically exist in the direction of maximum growth and calcification (Gladfelter et al., 1989; Taylor, 1977). The extent to which individual polyps might actively modulate their internal chemical milieu and affect their energetics spatially within a colony is an understudied area of coral biology.

Coral colonies are networks of individual polyps, and thus physiological models that consider the modular integration of corals are necessary for developing a robust understanding of colony physiology. While there are many ways to model a network as the field of network science is rapidly developing in biology and ecology (Proulx et al., 2005), we chose to use an electrical network model because it allows us to focus on the individual components before expanding the model to the whole colony. An electrical network models “flow” of something through an organism using circuit components such as resistors and capacitors. In the model, current is analogous to a flux of a metabolite, and voltage is analogous to the concentration of the metabolite. Electrical network modeling has been used to study physiological processes in organisms where modules interact. Nobel and Jordan (1983) developed an electrical network model to analyze transpiration rates and water potentials in three very morphologically different desert plants. Other studies used electrical network models for the vertebrate respiratory (Campbell and Brown, 1963) and circulatory systems (Dawson et al., 1982), excretory systems (Goldstein and Rypins, 1992), and the process of passive suspension feeding in lower invertebrates (Patterson, 1991). All of these electrical network modeling frameworks allow for testable predictions of physiological processes.

We define the time constant of mixing as the time needed for the gastrovascular system to mix and disperse most of the fluid in the coelenteron. We used the electrical network model to interpret microelectrode measurements of dissolved oxygen to calculate this time constant during a light-to-dark transition inside the coelentera of an imperforate coral, *Montastraea cavernosa*. Imperforate corals do not possess the ability of fluid exchange at the whole colony level, since individual polyps are only connected to their nearest neighbors, and only when the polyps are expanded. This approach allowed us to investigate whether there are one, or more, time scales affecting how mixing occurs inside coral polyps. Knowledge of these time constants provides insight into how quickly a polyp can respond to changing (internal or external) environmental conditions. Until now, these time scales have not been measured in scleractinians, but are important for understanding coral metabolism, including responses to global climate change.

## Materials and Methods

### Electrical network model

We constructed an electrical network model of gas flux in single polyps for perforate and imperforate corals. Our modeling framework makes testable predictions of dissolved oxygen concentration in the gastrovascular system (Fig. 1A and see Table 1 for parameter descriptions). In this framework, voltage is the oxygen concentration (the driving pressure) at any point in the system, and current is the change in oxygen concentration over time, i.e., the rate at which oxygen passes through a compartment in the system. Voltage sources represent external (environmental) and internal (photosynthesizing, symbiotic algae) sources of oxygen. Resistors reduce current flow in an electrical circuit. In our framework, they represent tissues, or water-filled volumes like the coelenteron, through which the current of oxygen must pass. Resistance is formalized as the time needed to pass oxygen through a model component on a per-volume basis. Capacitors are devices that store electrical energy. In our model, they represent volumes of compartments.

**Figure 1.**
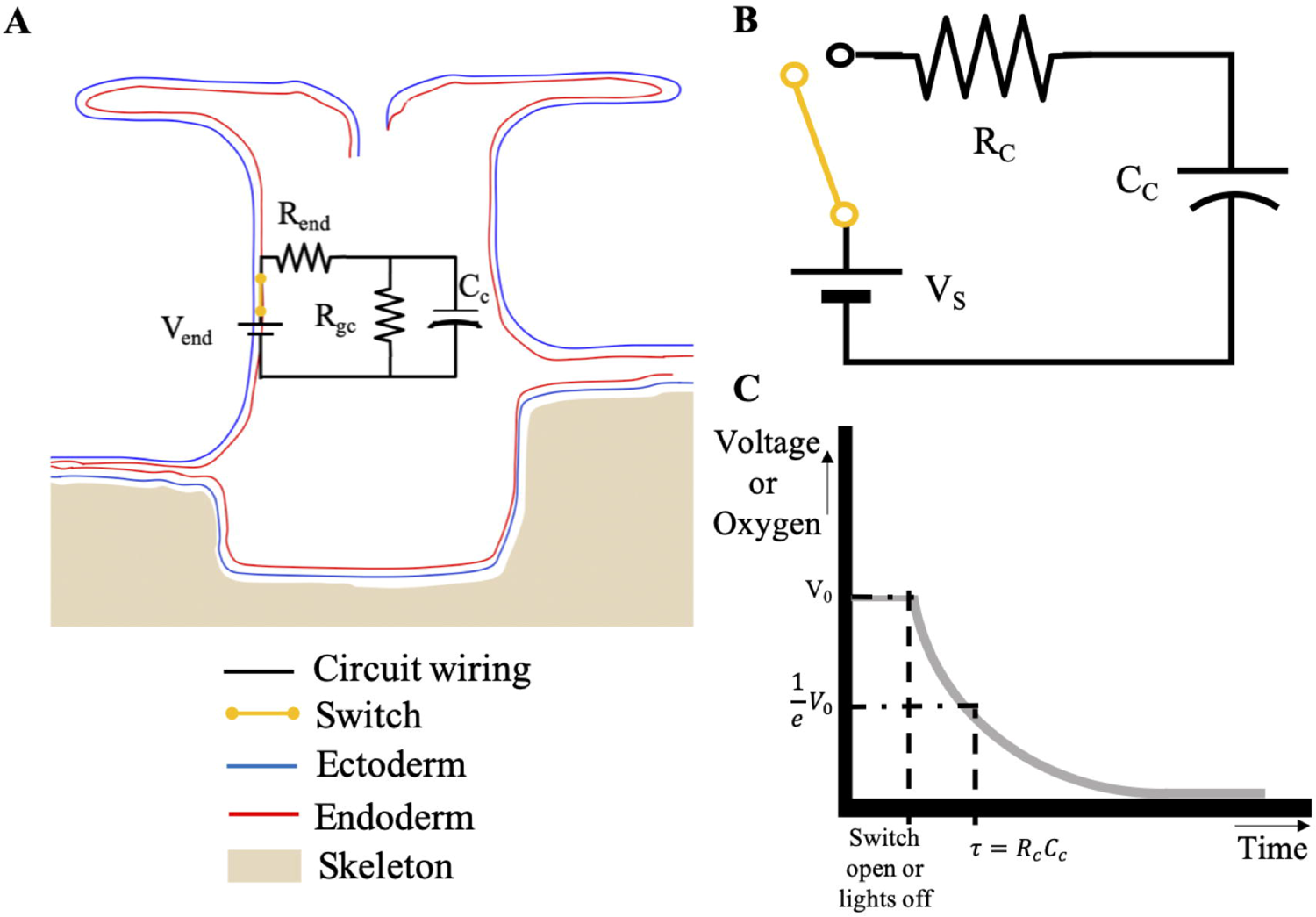
Conceptional diagram of the electrical network model of the scleractinian coelenteron and circuit dynamics. A) Circuit modeling oxygen dynamics in the coelenteron. See Table 1 for parameter explanations. B) Thévenin equivalent circuit of the scleractinian coelenteron circuit in A with the switch open representing the loss of the symbiont voltage source in the dark. C) Circuit (oxygen) dynamics when the switch is opened (light-to-dark transition) and the capacitor discharges across the resistor.

**Table 1.** Parameters in the electrical network model of the scleractinian coelenteron.

| Parameter | Circuit Component | Biological Meaning |
| --- | --- | --- |
| $V_s$ | Voltage source from symbionts | Algal symbionts in endoderm are an oxygen source when illuminated |
| $R_{end}$ | Resistor associated with endoderm | Resistance to flow of oxygen in the endoderm |
| $R_{gc}$ | Resistor associated with the gastric cavity | Resistance to flow of oxygen in the gastric cavity |
| $C_c$ | Capacitor associated with coelenteron | Oxygen storage capacity of the coelenteron |
| $R_c$ | Thevenin equivalent resistance for the coelenteron | Combined resistance to oxygen flow in the coelenteron from the endoderm and the gastric cavity |
| $V_c$ | Voltage across the coelenteron capacitor | Concentration of oxygen within the coelenteron |

Oxygen dynamics within the coelenteron are modeled as a simple resistor-capacitor circuit. Symbiotic algae in the endoderm serve as the main source of oxygen to the coelenteron (V_s_ in Fig. 1A and Table 1). There may also be some fluid, and thus oxygen, exchange with the environment through the mouth driven by ciliary movement of the siphonoglyphs (e.g., *Metridium senile;* Batham and Pantin, 1950). However, our electrical network model for the coelenteron assumes this environmental source of oxygen to be negligible compared to the oxygen source provided by the algal symbionts. There is a resistor, R_end_, associated with the endodermis as it acts as a barrier to the flux of oxygen into the coelenteron. R_gc_ represents the resistance to oxygen flux in the fluid of the gastric cavity. The capacity of the coelenteron to store oxygen is modeled by the capacitor C_c_. The circuit in Fig. 1A can be further reduced by using Thévenin’s Theorem, which states that it is possible to simplify any linear circuit to an equivalent circuit with just a single voltage source and series resistance connected to a load (Fig. 1B). In the Thévenin equivalent circuit, there is just one resistor, R_C_, that represents the total resistance to oxygen flux from the symbiont voltage source to the coelenteron where voltage is stored by the capacitor C_C_.

When the switch is opened, the voltage source is cut from the circuit and the charged capacitor discharges across the resistor. As the capacitor discharges, the voltage (V_C_) follows an exponential decay function

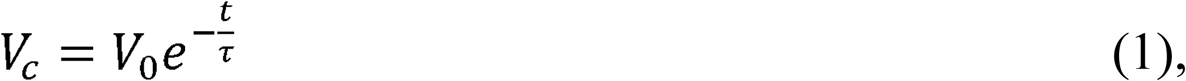

where *V*_0_ is the initial voltage (oxygen concentration) at the time (t) the switch is opened (Fig. 1C). The time constant (τ) is the time required to decrease the voltage to 36.8% of its initial value. Voltage dynamics when the switch is opened model the oxygen dynamics inside the coelenteron during a light-to-dark transition. When the lights are turned off, symbionts stop photosynthesizing and no longer provide a source of oxygen to the coelenteron. Levels of dissolved oxygen measured in the coral tissue are greater than air saturation level under light conditions but can drop to hypoxic levels within five minutes after a dark transition (Kühl et al., 1995). Fluid in the coelenteron is mixed by the cilia lining the endoderm and the oxygen is dispersed to the walls and mesenteries to be taken up during respiration. Thus, another way to view the time constant is the time it takes a polyp to mix and disperse 63.2% of the oxygen in the coelenteron to be consumed by tissues.

The time constant of mixing (τ) can be empirically determined by measuring the concentration of oxygen in the coelenteron during a light-to-dark transition and fitting the oxygen response curve with an exponential model. Once the time constant is known, the resistance associated with the coelenteron, R_C_, can be calculated by approximating the volume of the coelenteron, C_C_, as a cylinder, because τ = R_C_C_C_ (2).

### Oxygen measurements in Montastraea cavernosa

We measured the oxygen concentration within the coelentera of *M. cavernosa* polyps using a dissolved oxygen microelectrode (25-100 μm diameter tip) connected to a underwater amplifier and data logger (Unisense UWM-4, Aarhus Denmark, Fig. 2A). Microelectrodes allow for high spatial resolution and low analyte consumption. *M. cavernosa* has large polyps for easy insertion of microsensors, and is known to have two morphs, diurnal and nocturnal, with respect to expansion activity, with the nocturnal morph possessing larger polyps (Lasker 1977, 1981). Colonies of *M. cavernosa* were maintained in a flume for a controlled no-flow environment during measurement. Measurments were made in expanded polyps with clearly visible mouths. The microelectrode was positioned using a Narishige M-3333 micromanipulator (Narishige, Amityville, NY, USA). We inserted the microelectrode through the polyp mouth into the coelenteron (Fig. 2B and 2C). This process was visualized with the aid of a jewler’s magnifying headset. Once the signal had stabilized as shown on the Unisense data logger, the lights were turned off. Data were recorded continuously (1 Hz) until the signal stabilized to a new lower value, approximately ten minutes per trial. Lights were turned back on and the measurement procedure was repeated after the signal stabilized at the higher value once more, typically another ten minutes. We measured the change in oxygen concentration over time after a light-to-dark transition five times per polyp for three polyps each, in three different colonies of *M. cavernosa*. The three colonies were of comparable size, surface area of 106 - 150 cm^3^ as measured using image analysis (ImageJ; Schneider et al., 2012), and each had 176 to 242 polyps (Table 2).

**Figure 2.**
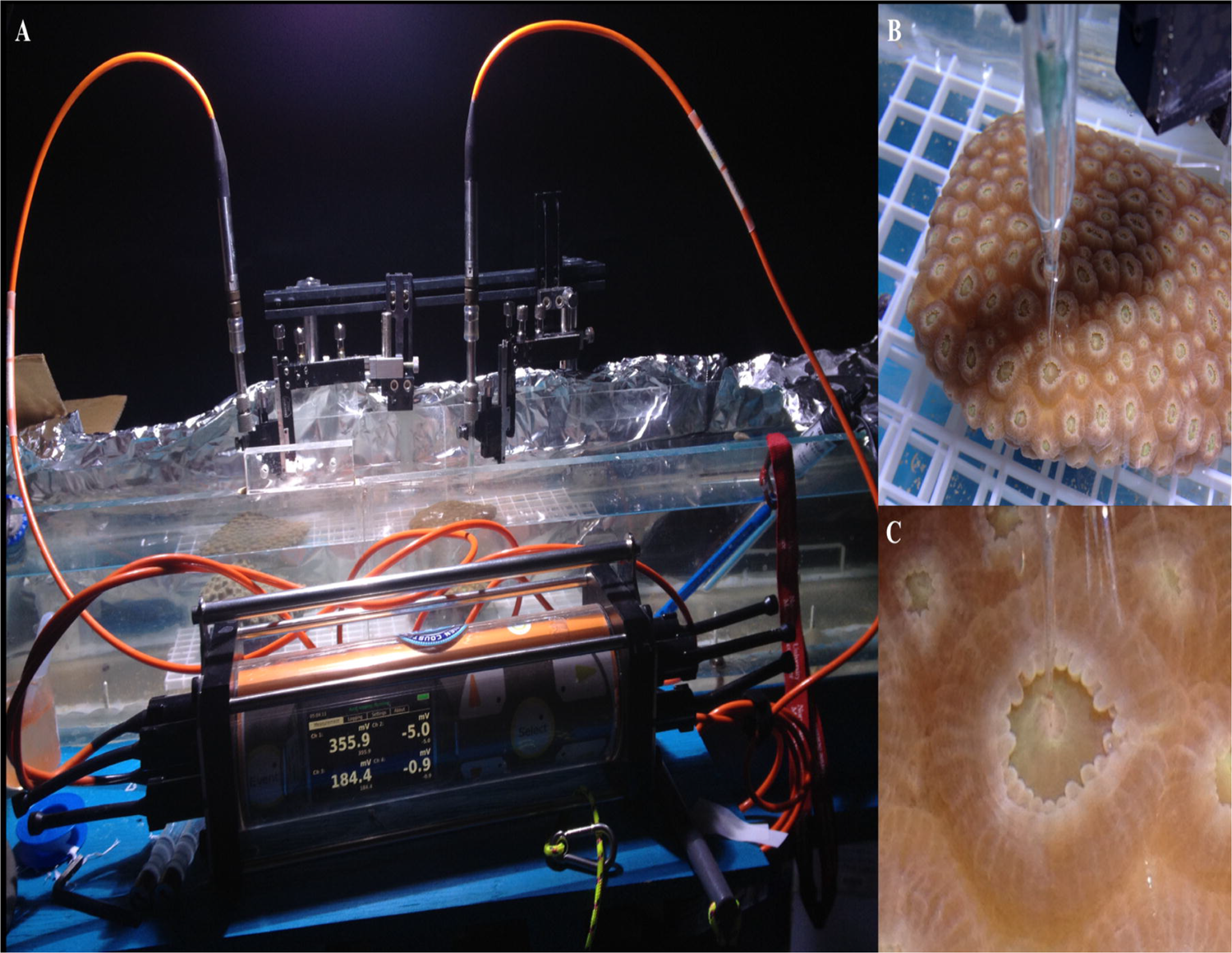
Experimental set up of oxygen measurements made within the coelenteron of *M. cavernosa* during a light-to-dark transition. A) Unisense underwater meter (UWM-4) with two oxygen microsensors and amplifiers positioned in the corals in a flume using the micromanipulators. B) Overhead view of the microsensor positioned in the coral. C) Closeup of the microsensor inserted through the mouth of the polyp.

**Table 2.** Weighted mean (± S.E.) results of the time constant of mixing and higher frequency oscillation analyses for all three colonies of *M. cavernosa*.Footnote: Approximate surface area was measured using ImageJ analysis of photos of the colonies next to a scale. Polyps were counted in ImageJ and the numbers are approximate because some polyps were hidden from the top-down photograph view. All means are weighted means ± S.E.

| Measure | Colony A | Colony B | Colony C | Species mean |
| --- | --- | --- | --- | --- |
| **Approx. surface area**<br>**(cm<sup>2</sup>)** | 110 | 150 | 106 | - |
| **Approx. number of**<br>**polyps (N)** | 176 | 242 | 184 | - |
| **Time constant of**<br>**mixing, $\tau$ (s)** | 113.4 ± 6.1 | 185.7 ± 34.6 | 187.7 ± 19.8 | 162.3 ± 24.4 |
| **Period of the**<br>**primary peak (s)** | 53.6 ± 2.3 | 46.9 ± 0.4 | 44.3 ± 1.8 | 48.3 ± 2.8 |
| **Power of primary**<br>**peak (ppm<sup>2</sup>)** | 1.4e<sup>-2</sup> ± 2.6e<sup>-2</sup> | 1.41e<sup>-2</sup> ± 4.1e<sup>-2</sup> | 1.6e<sup>-2</sup> ± 3.0e<sup>-2</sup> | 1.3e<sup>-2</sup> ± 0.2e<sup>-2</sup> |
| **Period of the**<br>**secondary peak (s)** | 32.6 ± 3.2 | 40.4 ± 4.0 | 34.8 ± 0.3 | 35.9 ± 2.3 |
| **Power of secondary**<br>**peak (ppm<sup>2</sup>)** | 5.7e<sup>-3</sup> ± 0.5e<sup>-3</sup> | 4.8e<sup>-3</sup> ± 1.0e<sup>-3</sup> | 9.3e<sup>-3</sup> ± 0.6e<sup>-3</sup> | 6.6e<sup>-3</sup> ± 1.4e<sup>-3</sup> |
| **Integrated power,**<br>**energy, (ppm<sup>2</sup>**<br>**radians s<sup>-1</sup>)** | 1.2e<sup>-3</sup> ± 1.9e<sup>-4</sup> | 0.7e<sup>-3</sup> ± 2.9e<sup>-4</sup> | 1.1e<sup>-3</sup> ± 3.0e<sup>-5</sup> | 1.0e<sup>-3</sup> ± 0.1e<sup>-3</sup> |
| **Ratio of $\tau$ to the**<br>**primary period** | 2.1 ± 0.1 | 4.1 ± 0.6 | 4.6 ± 0.4 | 3.6 ± 0.8 |

### Time constant of mixing analysis

All analyses were done in R v4.0.2 (R Core Team, 2017). Oxygen response curves were fit with an exponential model using the ‘vegan’ package (Oksanen et al., 2019; Fig. 3A). Time constants of mixing were then extracted from the fit equation. Time constants were log transformed to meet normality and homoscedasticity requirements before being fit with a linear mixed-effects model where polyp was set as a random effect. An ANOVA (Type I) was used to test for significant differences among the colonies. To test for significant differences among polyps in a single colony, data were fit with a linear model and again tested with an ANOVA (Type I).

**Figure 3:**
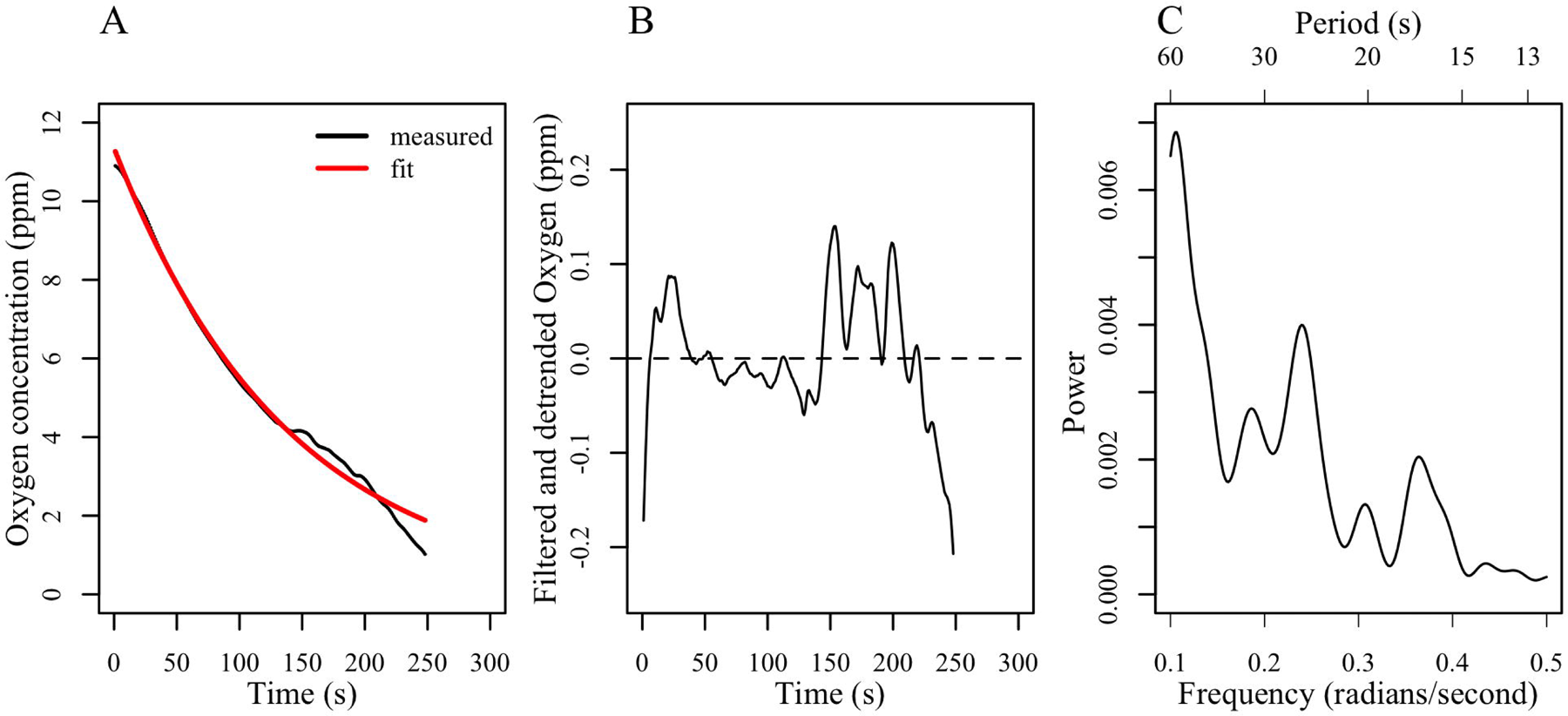
Steps of the fitting and spectral analysis procedure. A) Time series of dissolved oxygen measured using a microsensor inside a single polyp of *M. cavernosa*, with fitted exponential decay function (Eqn. 1) used for detrending the data. Note the very small oscillations discernible as the oxygen concentration is reduced to hypoxic levels. B) The detrended time series of dissolved oxygen. Series was detrended by subtracting the value of the exponential decay function at each time point. It was then high-pass filtered for oscillations occurring at periods faster than 62.8 s (= 0.016 Hz). C) Power spectral density for the detrended high-pass filtered time series.

### Spectral analysis of higher frequency variations

In addition to the overall exponential decline described above, inspection of the time series for dissolved oxygen showed interesting higher frequency variations of low amplitude (c. 0.1 ppm), that often became more apparent as levels inside the polyp dropped to levels defined as hypoxia (< 2 ppm). To quantify the time scale of these variations, we computed the power spectral density function (ver. 12.1, Wolfram Research, https://www.wolfram.com). First, data were fit to an exponentially decaying function identical to eqn1">Eqn/xref> (Fig. 3A). This fitted function was evaluated for each point in the time series and was then subtracted from the corresponding datum for that time to produce a detrended time series (Fig. 3B). A power spectral density function with a Hann smoothing window to eliminate artifacts that arise from a finite dataset was applied to the detrended time series with Fourier coefficients set to those recommended for analysis of time series data (Fig. 3C). The resulting spectrum was high-pass filtered to pass only frequencies higher than 0.1 radians s^−1^, equivalent to a frequency of 0.016 Hz, or a period faster than 62.8 s (Fig. 3C). The power and period of the highest and second highest peaks present in the power spectral density function were then recorded for each trial. The primary peak, the frequency (period) with the highest power, had the most biological significance and represented a time scale for higher frequency oscillations in oxygen concentrations within the coelenteron.

Differences between the power (magnitude) and periods (time scales) of the two peaks were tested using Mann-Whitney tests. Period data were non-normal, so significant differences in the periods of the primary and secondary maxima among colonies were tested using 1000 Monte

Carlo simulations of a general linear mixed-effects model where the ANOVA (Type I) test statistic distribution was compared with the actual statistic. The powers of the spectral peaks were normally distributed and were analyzed using the same approach described in the previous section for the time constant of mixing (τ).

We also determined the integrated power spectral density for each trial over the frequency range of 0.1 - 0.5 radians s^−1^. The integrated power (ppm^2^ radians s^−1^) is a measure of the energy dissipated – the “churn” – within the coelenteron. Significant differences in integrated power were determined following the same statistical analysis used for the time constant of mixing.

## Results

### Model validation and the time scale of mixing

Oxygen concentration inside *M. cavernosa* coelentera during the light-to-dark transition followed the expected exponential decay function predicted by Eqn. 1. An example of the oxygen decay curves for all five trials in one polyp is given by Fig. 4. The coefficient of determination values (R^2^ values for each trial presented in Supplement S1) ranged from 0.69 to 1.00 with an average value of 0.96, indicating that Eqn. 1 explained most variation in the data. In addition to the overall exponential decay, upon inspection and as described above, the oxygen response curves showed low-amplitude, higher frequency variations as oxygen inside the polyp dropped below hypoxic levels. These low-amplitude oscillations indicate that the coelenteron has behavior not predicted by the framework of the electrical network model. As discussed later, these low-amplitude, high-frequency oscillations give a time scale for mixing of the coelenteron that may represent exchange with the environment and/or within the gastrovascular system.

**Figure 4:**
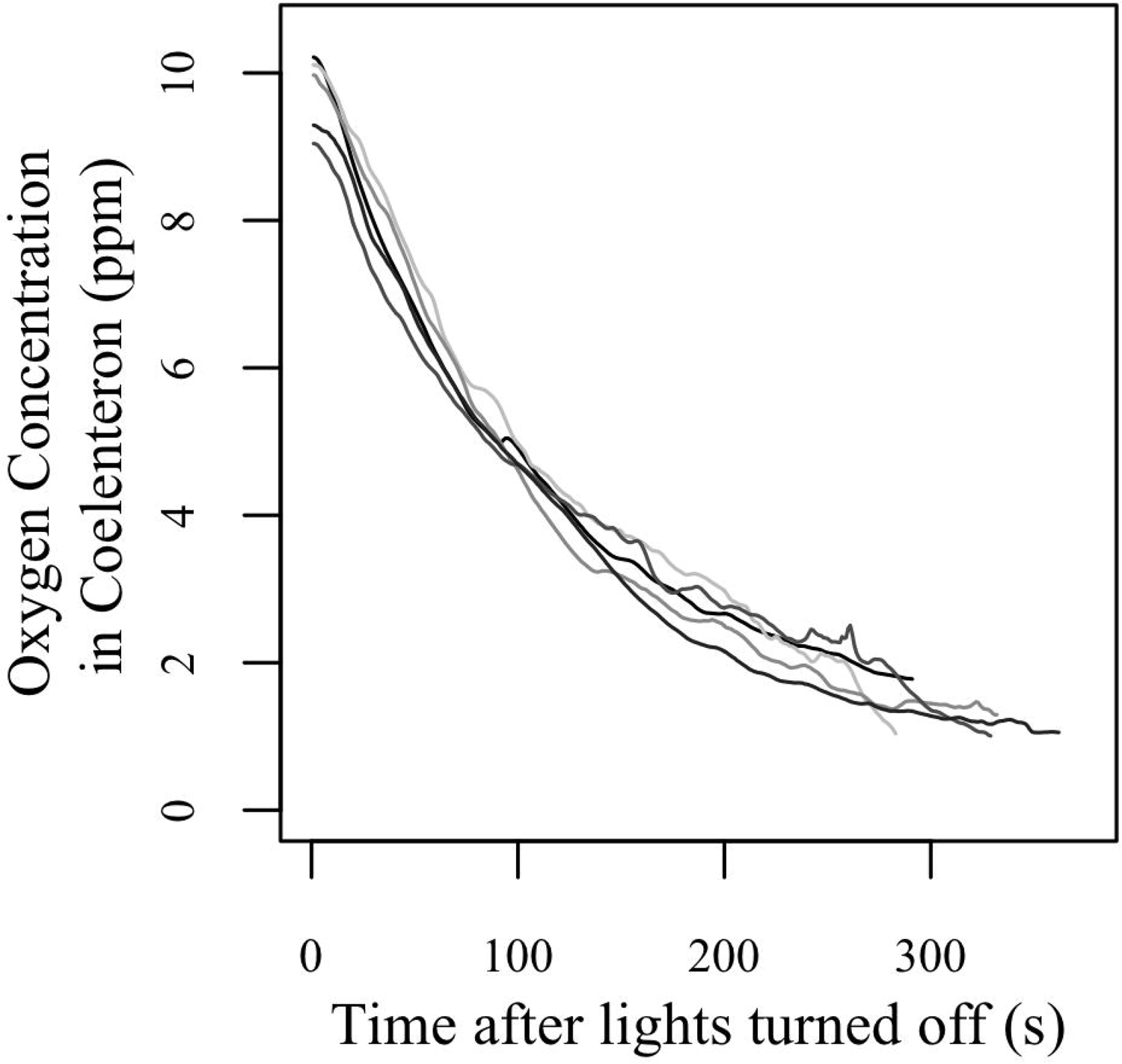
Time series of dissolved oxygen concentrations during a light-to-dark transition for all trials (*n* = 5) in a polyp of colony C. Each trial is a separate line.

The dominant time scale of mixing for *M. cavernosa* was on the order of 70.8 to 371.9 seconds (roughly 1-6 minutes, Fig. 5). On average, the species had a time constant of mixing of about three minutes (162.3 ± 24.4 s, mean ± S.E., Table 2). The three colonies tested did not have significantly different time constants of mixing (*P* = 0.09). Colony A had the shortest time scale of mixing with τ = 113.4 ± 6.1 s, but colonies B (τ = 185.7 ± 34.6 s) and C (τ =187.7 ± 19.8 s) had time constants on the order of a minute longer. Within each colony, the time constants varied slightly across polyps; however no significant differences were found (*P* > 0.05, Fig. 5).

**Figure 5.**
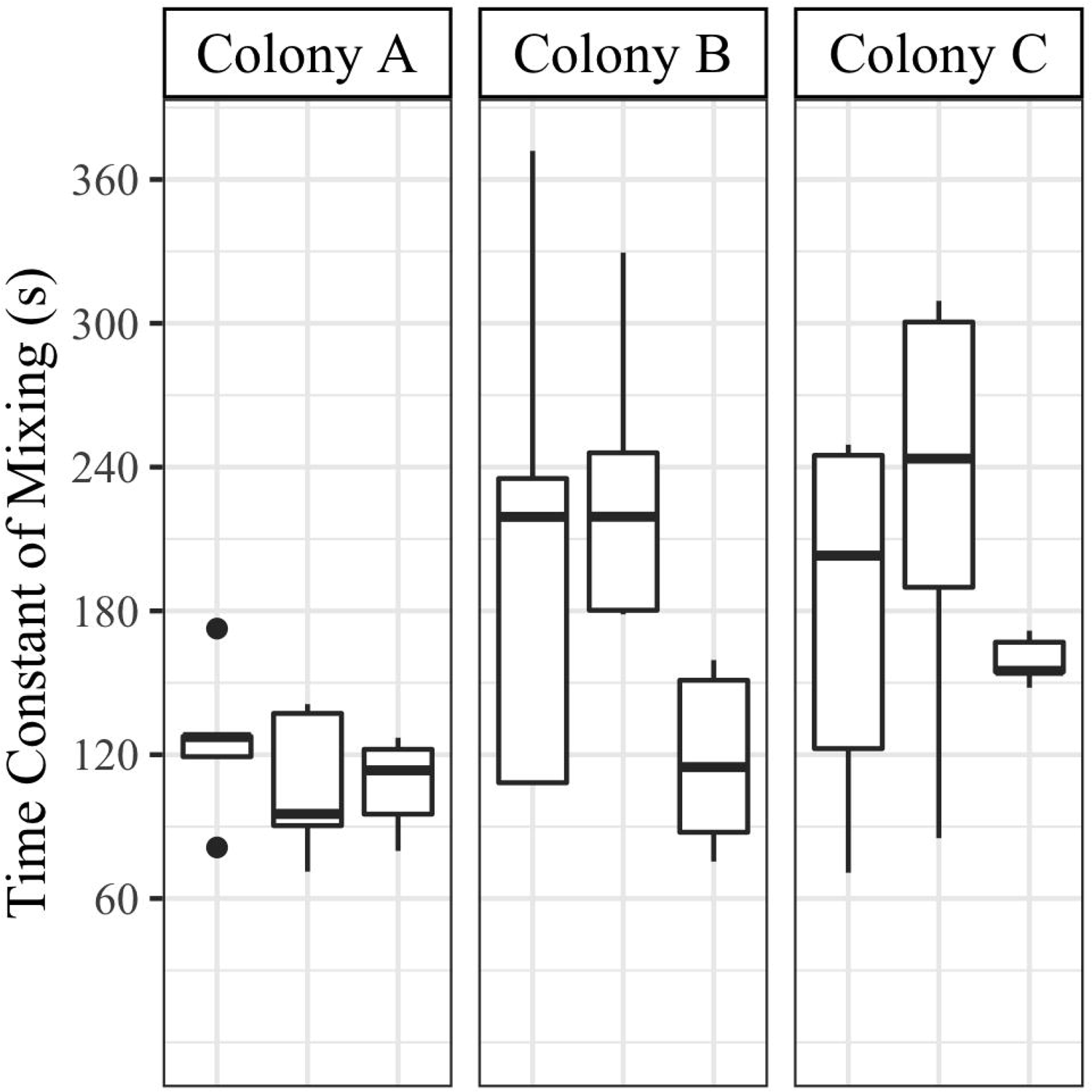
Time constant of mixing (τ) measurements in three polyps of three colonies of *M. cavernosa*. The three colonies did not have significantly different time constants (ANVOA Type I, *P* = 0.09). However, there is a small amount of individual variation of τ among and within colonies. τ of polyps within a colony did not significantly vary for all colonies (ANOVA Type I, *P* > 0.05). The figure shows the median, first and third quartiles, largest value within 1.5 times the interquartile range above the 75^th^ percentile, and the smallest values within 1.5 times the interquartile range below the 25^th^ percentile. Outliers are outside the 1.5X the interquartile range beyond either ends. *n* = 5 for each polyp.

### Resistance to oxygen flow in the coelenteron

From the electrical network model, we estimated the resistance to oxygen flow in the coelenteron using the determined time constant of mixing, as 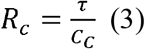 (3). We approximated the polyp as a cylinder and estimated the internal volume (capacitance, C_c_). The radius of a *M. cavernosa* polyp is approximately 0.5 cm and if we consider an expanded polyp height of 1 cm, the volume is approximately 0.8 cm^3^. Using the species average τ = 162.3 s, the resistance, R_c_, was computed to be ~ 203 s cm^−3^. Resistance to oxygen flow will vary with polyp size and expansion state.

### Higher frequency variations in oxygen concentration

Periods of the primary and secondary maxima of the higher frequency oscillations in oxygen concentration were determined to be on the order of every 20–60 seconds (Table 2; Supplement S1). The two periods were significantly different (*P* < 0.001), thus indicating that there are at least two time scales of higher frequency mixing occurring in the coelenteron. The frequency with the highest power, the primary peak, revealed that there were small oscillations (~ 0.1 ppm) in oxygen concentration occurring every 48.3 (mean) ± 2.8 (S.E.) seconds in the coelenteron of *M. cavernosa*. A secondary peak with a lower power suggested that there was a second oscillatory rhythm occurring every 35.9 ± 2.3 seconds on average in *M. cavernosa*. There were no significant differences in primary (*P* = 0.091) and secondary (*P* = 0.128) peak period values among the three colonies tested.

Power (ppm^2^), or magnitudes, of the two largest peaks were significantly different (*P* < 0.001). These powers represent the strength of the fluctuations at the two periods (Table 2). The power of the primary peak was significantly higher than that of the secondary peak, and it did not vary across the colonies measured (*P* = 0.13). Power of the secondary peak did not vary across colony either (*P* = 0.28).

Integrated power (ppm^2^ radians s^−1^), a measure of the energy dissipated over the entire trace, varied significantly across all colonies (*P* = 0.04). The species average for integrated power is 1.0e^−3^ ± 0.1e^−3^ ppm^2^ radians s^−^1. A *post hoc* comparison test of the integrated power (energy) found that colony B had an integrated power significantly lower than colonies A and C (*P* = 0.01 and *P* = 0.006 respectively).

Additionally, we calculated the ratio of the time constant of mixing (τ) to the period of the primary peak (Table 2). This ratio represents the relationship between the dominant time scale of mixing and the higher frequency time scales of mixing in the coelenteron. For the *M. cavernosa* colonies measured, the average of this dimensionless ratio was 3.6 ± 0.8. The dominant time scale of mixing was approximately 3-4 times longer than the high frequency time scales in the coelenteron. This ratio varied significantly across the colonies measured (*P* = 0.002). Colony A had the lowest value (2.1 ± 0.1), while colonies B (4.1 ± 0.6) and C (4.6 ± 0.4) were similar and twice the magnitude of A.

## Discussion

We explored time scales of mixing in *M. cavernosa* using a novel electrical network model of oxygen dynamics in the scleractinian gastric cavity and *ex situ* microelectrode measurements of coelenteric dissolved oxygen concentrations in a laboratory mesocosm. Our metric, the time constant of mixing (τ) captures the dominant time scale of mixing in the coelenteron, the time it takes a polyp to mix and disperse 63.2% of the dissolved oxygen in the coelenteron. The time constant of mixing was on the order of three minutes and did not vary significantly across colonies of *M. cavernosa*. This time scale is much faster than initial predictions based solely on diffusion processes. Our electrical network model accurately predicted the overall oxygen dynamics in *M. cavernosa* coelentera during a light-to-dark transition; however, we also found small (~0.1 ppm) oscillations in dissolved oxygen concentration. The two higher frequency time scales of mixing were on the order of 50 s and 35 s. Mixing in the coelenteron, the largest component by volume in the scleractinian gastrovascular system, occurs at multiple time scales all as fast or faster than environmental conditions can change. The most rapid changes seen routinely on coral reefs are perturbations of water chemistry caused by internal waves, and these have time scales on the order of minutes (Wolanksi and Delesalle, 1995; Leichter et al., 1996; Wall et al., 2015).

What could account for the faster time scales of mixing? Possibilities that could explain these rhythms include 1) water brought into the polyp’s coelenteron from neighbors via the imperforate connections, 2) exchange with the surrounding seawater modulated by the polyp’s pharynx, and 3) subtle muscular contractions that change the spacing of the mesenteries relative to the polyp wall allowing for pockets of more oxygenated fluid to mix in with the rest of the gastric fluid. We searched for episodic exchange between *M. cavernosa* polyps using sodium fluorescein dye mixed with seawater and a UV light following the methods by Gladfelter (1983), but did not observe impulsive exchange because the coral’s own natural fluorescence interferes with dye visualization. We did not test for intake of water episodically through the pharynx because Gladfelter (1983) never observed this behavior in *A. cervicornis*. During microsensor studies in other species, researchers have noted that the tissue around a polyp mouth frequently contracted and expanded (e.g. Kühl et al., 1995); however, we did not observe obvious muscular contractions of the polyp during our measurements.

The higher frequency oscillations of dissolved oxygen that we observed in *M. cavernosa* may be triggered by hypoxia. Hypoxia is more likely to occur at night on coral reefs from community respiration, although daytime hypoxic conditions are becoming more frequent on some reef systems (Nelson and Altieri, 2019). At night when polyps are expanded, if connections are patent between polyps, the oscillations we observe may be an attempt to regulate oxygen upward if a polyp becomes stressed. A perforate species like *A. cervicornis*, that has a more integrated gastrovascular system may have a shorter dominant time scale of mixing along with more variable high frequency time scales. Whether whole colony integration affects these time scales of mixing should be tested to further explore interspecific difference in responses to changing environmental conditions, like hypoxia.

Colony and individual polyp size may play a role in internal mixing dynamics. Colony B was our largest colony, both in surface area and number of polyps. We found no significant differences in the time scales of mixing for B and the other two colonies; however colony B had a significantly lower integrated power (energy) than the other two colonies. The integrated power is a measure of the energy dissipated over the total time that we measured dissolved oxygen, i.e., a measure of the internal churn associated with mixing and dispersing dissolved oxygen in the coelenteric fluid. The largest coral had the lowest amount of internal churn possibly indicating that size plays a role in the magnitude of high frequency mixing in coral coelentera. Additionally, the smallest colony (A) had a significantly lower ratio of the time constant of mixing to the primary period of higher frequency mixing. The dominant time scale of mixing is only ~ 2X longer than the time scale of higher frequency mixing in colony A, whereas it is ~ 4X longer in the other two, larger colonies.

Colony level differences may also be intraspecific differences in metabolism related to colony genetics similar to that observed for bleaching response to temperature increase (Morikawa and Palumbi, 2019). Colony shape and flow regime have a direct effect on external mass transfer dynamics and local metabolism (Patterson, 1992a,b), and thus we may expect to find differences in internal mixing with variable colony shapes, sizes, polyp expansion, and environmental flow regimes. Time scales of mixing should be measured in polyps from a range of colony sizes in order to further explore a potential size-dependent relationship of internal mixing in scleractinians.

We did not make explicit measurements of polyp heights (i.e., extension levels) while recording the oxygen dynamics, as we only ensured that polyps stayed expanded during measurements. Polyp expansion may prove to be an important variable affecting mixing, as the internal fluid volume will change with the state of polyp expansion. As the volume of the coelenteron increases, the surface area of the ciliated mesenteries increases as the second power of height or radius. These ciliated mesenteries are the primary drivers of fluid mixing in the gastrovascular system. There could exist an inherent limit to polyp size due to the ability of the coelenteron to mix fluid. Patterson (1992a,b) predicted polyp geometric dimensions to be diffusively similar, whereby polyps should change their diameter to height ratio in such a way as to keep the normalized diffusive flux into the polyp the same, but his theory has never been fully tested. Surveys of the time constant of mixing in relation to varying polyp expansion levels and geometries will provide an avenue to test Patterson’s (1992a,b) theory.

Our measurements were made under controlled, no-flow conditions. It would be interesting to test whether environmental flow affects internal mixing and polyp behavior since it is known to affect external mass transfer dynamics (Patterson, 1992a,b). Furthermore, internal mixing will drive mass transfer dynamics in the coelenteron. Environmental flow has been shown to increase photosynthetic performance in corals by increasing the oxygen efflux from the organism to the water (Mass et al., 2010), and thus we could also consider the role of fluid mixing and residence time in the gastric cavity as a potential regulator for photosynthetic productivity. Our measurement protocol could not capture the full mass transfer dynamics of the internal environment because we measured the oxygen dynamics at a single point. Although this allowed us to measure previously unobserved time scales of mixing in the coelenteron, we may be missing other important dynamics. Advances in sensor technology armed with this new knowledge of the underlying time scales of mixing may now be able to elucidate new insights into mass transfer dynamics inside of the coral gastrovascular system.

Although digestion in anthozoans is understudied, most of the digestion of planktonic prey and organic matter takes place in the coelenteron where larger particles are first broken down by a digestive fluid into smaller particles that can be ingested by the endoderm cells (reviewed by Goldberg, 2018; Boschma 1925; Murdock 1978; Pratt, 1905). Nicol (1959) determined digestion times in a species of sea anemone to be on the order of 8–24 hours. We determined that mixing within a scleractinian polyp’s coelenteron is on the order of ~three minutes. This implies that the bioreactor of the coelenteron is well mixed *sensu* Penry and Jumars (1986), but this time scale of mixing will be important for nutrient transport within single polyps and throughout the gastrovascular system.

Future work to elucidate the mechanisms of mixing in scleractinian polyp coelentera and the gastrovascular system could use optical coherence tomography (Wangpraseurt et al., 2017) to examine whether tissues in the polyp move at the same period as the oscillations we detected. Further, magnetic nanoparticles, such as those used in vascular studies (Nacev et al., 2010), could be injected into the gastrovascular system, or in the boundary layer of water over the polyp, where they could be tracked to test whether water is entrained from adjacent polyps with the observed periodicity.

The residence time of water in a coral colony’s gastrovascular system will have an effect on the supply of ions for calcification and photosynthesis and will be a critical point of limitation for these processes. We determined that time scales of mixing in the coelentera of corals are on the order of three minutes or faster. These time scales are fast enough to keep up with changes in variable environmental conditions on the reef within a diel cycle (Levy et al., 2006), due to internal waves (Leichter et al., 1996), and upwelling (Wolanski and Delesalle, 1995). If time scales of mixing are faster in perforate corals that have a highly integrated gastrovascular system, will these corals be more or less susceptible to changing environmental conditions? A meta-analysis by Swain et al. (2018) found that highly integrated coral species have a significantly reduced bleaching response. Bove et al. (2020) determined species-specific differences in response to ocean acidification when measuring pH levels in the coelenterons of two scleractinian species in relation to calcification. Interspecific variation in the time scales of mixing may provide new insight into how corals respond to stressors. More work needs to be done to understand the effects of gastrovascular architecture on internal mixing in corals if we are to make accurate predictions about physiological performance under global change (Morikawa and Palumbi, 2019).

## Acknowledgements

We thank E. Gladfelter and L. Carpenter for insights on coral gastrovascular physiology and methods discussion during the early phases of this research. B. Helmuth, T. Gouhier, E. Muller, S. Vollmer, A. Dwyer, and L. Allen-Jacobson provided helpful discussion. This material is based upon work supported by the National Science Foundation under Grant No. IOS-1146056 & IOS-1412462 (MRP), and Northeastern University. Any opinions, findings, and conclusions or recommendations expressed in this material are those of the authors and do not necessarily reflect the views of the National Science Foundation. SDW was supported by a National Science Foundation Graduate Research Fellowship. This is contribution XXX from the Marine Science Center, Northeastern University.

## Data Availability

Data and R code will be made available at https://github.com/saradwms/McavPolypMixing upon publication.

## Literature cited

Agostini, S., Suzuki, Y., Higuchi, T., Casareto, B. E., Yoshinaga, K., Nakano, Y., & Fujimura, H. 2012. Biological and chemical characteristics of the coral gastric cavity. Coral Reefs 31(1), 147–156.

Batham, E. J., and Pantin, C. F. A. 1950. Muscular and hydrostatic action in the sea-anemone *Metridium senile (L.)*. J. Exp. Biol. 27(3), 264–289.

Blackstone, N. W. 1996. Gastrovascular flow and colony development in two colonial hydroids. Biol. Bull. 190(1), 56–68.

Boschma, H. 1925. On the feeding reactions and digestion in the coral polyp *Astrangia danae*, with notes on its symbiosis with zooxanthellae. Biol. Bull. 49(6), 407–439.

Bove, C.B., Whitehead, R.F. & Szmant, A.M. 2020. Responses of coral gastrovascular cavity pH during light and dark incubations to reduced seawater pH suggest species-specific responses to the effects of ocean acidification on calcification. Coral Reefs 1–17.

Cai, W. J., Ma, Y., Hopkinson, B. M., Grottoli, A. G., Warner, M. E., Ding, Q., Hu, X., Yau, X., Schoepf, V., Xu, H., et al. 2016. Microelectrode characterization of coral daytime interior pH and carbonate chemistry. Nat. commun. 7(1), 1–8.

Campbell, D. and Brown, J. 1963. The electrical analogue of lung. Br. J. Anaesth. 35(11), 684–92.

Carpenter, L.W., and M.R. Patterson. 2007. Water flow influences the distribution of photosynthetic efficiency within colonies of the scleractinian *Montastrea annularis* (Ellis and Solander 1786): implications for coral bleaching. J. Exp. Mar. Biol. Ecol. 351, 10–26.

Carpenter, L.W., M.R. Patterson, and E.S. Bromage. 2010. Water flow influences the spatiotemporal distribution of heat shock protein 70 within colonies of the scleractinian coral *Montastrea annularis* following heat stress (Ellis and Solander, 1786): implications for coral bleaching. J. Exp. Mar. Biol. Ecol. 387: 52–59.

Comeau, S., Carpenter, R. C., and Edmunds, P. J. 2013. Coral reef calcifiers buffer their response to ocean acidification using both bicarbonate and carbonate. Proc. R. Soc. Lond., B, Biol. Sci. 280(1753), 20122374.

Dawson, C.A., J.H. Linehan, and D.A. Rickaby. 1982. Pulmonary microcirculatory thermodynamics. Ann. N. Y. Acad. Sci. 384, 90–106.

Duerden, J. E. 1902. West Indian madreporarian polyps (Vol. 8). US Government Printing Office.

Gateno, D., A. Israel, Y. Barki, and B. Rinkevich. 1998. Gastrovascular circulation in an octocoral: evidence of significant transport of coral and symbiont cells. Biol. Bull. 194(2), 178–186.

Gladfelter, E.H. 1983. Circulation of fluids in the gastrovascular system of the reef coral *Acropora cervicornis*. Biol. Bull. 165, 619–636.

Gladfelter, E. H., Michel, G., & Sanfelici, A. 1989. Metabolic gradients along a branch of the reef coral *Acropora palmata*. Bull. Mar. Sci. 44(3), 1166–1173.

Goldberg, W.M. 2018. Coral food, feeding, nutrition, and secretion: a review. In: Kloc M., Kubiak J. (eds) Marine Organisms as Model Systems in Biology and Medicine (ed. M. Kloc and J.Z. Kubiak). Chap, Switzerland: Springer, pp. 377–421.

Goldstein, L.J., and E.B. Rypins. 1992. A computer model of the kidney. Comp. Meth. Prog. Biomed. 37(3), 191–203.

Harmata, Katherine L., Austin P. Parrin, Patrick R. Morrison, Kristin K. McConnell, Lori S. Bross, and Neil W. Blackstone. 2013. Quantitative measures of gastrovascular flow in octocorals and hydroids: toward a comparative biology of transport systems in cnidarians. Invertebr. Biol. 132(4), 291–304.

Jokiel, P. L. (2011). The reef coral two compartment proton flux model: A new approach relating tissue-level physiological processes to gross corallum morphology. J. Exp. Mar. Biol. Ecol., 409(1-2), 1–12.

Jones, W.C., V.J. Pickthall, and S.P. Nesbitt. (1977). A respiratory rhythm in sea anemones. J. Exp. Biol. 68, 187–198.

Kühl, M., Cohen, Y., Dalsgaard, T., Jørgensen, B. B., and Revsbech, N. P. (1995). Microenvironment and photosynthesis of zooxanthellae in scleractinian corals studied with microsensors for O2, pH and light. Mar. Ecol. Prog. Ser. 117(1-3), 159–172.

Lasker, H.R. (1977). Patterns of zooxanthellae distribution and polyp expansion in the reef coral *Montastrea cavernosa*. Proc. 3rd Int’l Coral Reef Symp. 607–613.

Lasker, H.R. (1981). Phenotypic variation in the coral *Montastrea cavernosa* and its effect on colony energetics. Biol. Bull. 160, 292–302.

Leichter, J.J., S.R. Wing, S.L. Miller, and M.W. Denny. (1996). Pulsed delivery of subthermocline water to Conch Reef (Florida Keys) by internal tidal bores. Limnol. Occeanogr. 41, 1490–1501.

Levy, O, Y. Achituv, Y.Z. Yacobi, Z. Dubinsky, and N. Stambler. (2006). Diel ‘tuning’ of coral metabolism: physiological response to light cues. J. Exp. Biol. 209, 273–283.

Mass, T., Genin, A., Shavit, U., Grinstein, M., and Tchernov, D. (2010). Flow enhances photosynthesis in marine benthic autotrophs by increasing the efflux of oxygen from the organism to the water. Proc. Natl. Acad. Sci. USA. 107, 2527–2531.

Morikawa, M.K., and S.R. Palumbi. (2019). Using naturally occurring climate resilient corals to construct bleaching-resistant nurseries. Proc. Natl. Acad. Sci. USA. 116(21), 10586–10591.

Murdock, G. R. (1978). Circulation and digestion of food in the gastrovascular system of gorgonian octocorals (Cnidaria; Anthozoa). Bull. Mar. Sci. 28(2), 363–370.

Nacev, A., C. Beni, O. Bruno, and B. Shapiro. (2010). Magnetic nanoparticle transport within flowing blood and into surrounding tissue. Nanomedicine (Lond.) 5(9), 1459–1466.

Nelson, H.R., and Altieri, A.H. (2019). Oxygen: the universal currency on coral reefs. Coral Reefs. 38, 177–198.

Nicol, J.A.C. (1959). Digestion in sea anemones. J. Mar. Biolog. Assoc. U.K. 38, 469–477.

Nobel, P.S., and Jordan, P.W. (1983). Transpiration stream of desert species: resistances and capacitances for a C3, a C4, and a CAM plant. J. Exp. Bot. 34, 1379–1391.

Jari Oksanen, F. Guillaume Blanchet, Michael Friendly, Roeland Kindt, Pierre Legendre, Dan McGlinn, Peter R. Minchin, R. B. O’Hara, Gavin L. Simpson, Peter Solymos, M. Henry H. Stevens, Eduard Szoecs, and Helene Wagner. (2019). vegan: Community Ecology Package. R package version 2.5-5.

Parrin, A. P., Netherton, S. E., Bross, L. S., McFadden, C. S., and Blackstone, N. W. (2010). Circulation of fluids in the gastrovascular system of a stoloniferan octocoral. Biol. Bull. 219(2), 112–121.

Patterson, M.R. (1991). Passive suspension feeding by an octocoral in plankton patches: empirical test of a mathematical model. Biol. Bull. 180(1), 81–92.

Patterson, M. R. (1992a). A mass transfer explanation of metabolic scaling relations in some aquatic invertebrates and algae. Science. 255(5050), 1421–1423.

Patterson, M.R. (1992b). A chemical engineering view of cnidarian symbioses. Am. Zool. 32(4), 566–582.

Patterson, M. R., Sebens, K. P., and Olson, R. R. (1991). *In situ* measurements of flow effects on primary production and dark respiration in reef corals. Limnol. Oceanog. 36(5), 936–948.

Penry, D.L., and P.A. Jumars. (1986). Chemical reactor analysis and optimal digestion. BioScience. 36(5), 310–315.

Pratt, E. M. (1905). The digestive organs of the Alcyonaria and their relation to the mesogloeal cell plexus. Quart. J. Micr. Sci. 49, 327–362.

Proulx, S. R., Promislow, D. E., and Phillips, P. C. (2005). Network thinking in ecology and evolution. Trends Ecol. Evol. 20(6), 345–353.

R Core Team (2017). R: A language and environment for statistical computing. R Foundation for Statistical Computing, Vienna, Austria. https://www.R-project.org/.

Schneider, C. A.; Rasband, W. S. and Eliceiri, K. W. (2012). NIH Image to ImageJ: 25 years of image analysis. Nat. Methods 9(7), 671–675.

Swain, T. D., Bold, E. C., Osborn, P. C., Baird, A. H., Westneat, M. W., Backman, V., & Marcelino, L. A. (2018). Physiological integration of coral colonies is correlated with bleaching resistance. Mar. Ecol. Prog. Ser. 586, 1–10.

Taylor, D. L. (1977). Intra-colonial transport of organic compounds and calcium in some Atlantic reef corals. Proc. 3rd Int’l Coral Reef Symp. 431–436.

Wall, M., L. Putchim, G.M. Schmidt, C. Jantzen, S. Khokiattiwong, and C. Richter. (2015) Large-amplitude internal waves benefit corals during thermal stress. Proc. R. Soc. B. 282, 20140650.

Wangpraseurt, D., C. Wentzel, S.L. Jacques, M. Wagner, and M. Kühl. (2017). *In vivo* imaging of coral tissue and skeleton with optical coherence tomography. J. R. Soc. Interface 128, 20161003.

Wolanski, E., and Delesalle, B. (1995). Upwelling by internal waves, Tahiti, French Polynesia. Cont. Shelf. Res. 15(2-3), 357–368.

