## Supplemental Table 1 for "Time scales of mixing in an imperforate scleractinian coelenteron"

| *Col.* | *Pol.* | *Exp.* | *$R^2$* | *τ (s)* | *PP Per. (s)* | *PP Pow. ($ppm^2$)* | *SP Per. (s)* | *SP Pow. ($ppm^2$)* | *N* | *IP ($ppm^2$ radians $s^{-1}$)* | *τ: PP* |
|---|---|---|---|---|---|---|---|---|---|---|---|
| *A* | A1 | -0.012 | 0.980 | 81.301 | 54.2 | 4.63E-03 | 31.1 | 1.90E-03 | 107 | 4.72E-04 | 1.50 |
| *A* | A1 | -0.008 | 0.863 | 119.175 | 61 | 2.17E-02 | 32.1 | 1.34E-02 | 211 | 1.53E-03 | 1.95 |
| *A* | A1 | -0.008 | 0.956 | 127.361 | 59.1 | 6.87E-03 | 26.2 | 4.01E-03 | 248 | 7.35E-04 | 2.16 |
| *A* | A1 | -0.008 | 0.966 | 127.651 | 62.8 | 2.27E-02 | 38.3 | 4.65E-03 | 279 | 8.16E-04 | 2.03 |
| *A* | A1 | -0.006 | 0.974 | 172.618 | 46.5 | 8.49E-03 | 29.4 | 4.42E-03 | 340 | 5.70E-04 | 3.71 |
| *A* | A2 | -0.014 | 0.991 | 71.205 | 34 | 1.38E-02 | 24.6 | 4.69E-03 | 158 | 1.83E-03 | 2.09 |
| *A* | A2 | -0.010 | 0.988 | 95.289 | 62.8 | 1.10E-02 | 29.8 | 1.03E-02 | 215 | 1.77E-03 | 1.52 |
| *A* | A2 | -0.011 | 0.996 | 90.402 | 36.5 | 2.19E-02 | 19.8 | 3.33E-03 | 204 | 1.98E-03 | 2.48 |
| *A* | A2 | -0.007 | 0.997 | 137.229 | 56.6 | 6.47E-03 | 35.7 | 3.75E-03 | 320 | 5.53E-04 | 2.42 |
| *A* | A2 | -0.007 | 0.990 | 141.103 | 55.6 | 1.68E-03 | 28.4 | 1.36E-03 | 308 | 2.19E-04 | 2.54 |
| *A* | A3 | -0.013 | 0.942 | 79.841 | 62.8 | 2.27E-02 | 40 | 2.52E-03 | 139 | 9.07E-04 | 1.27 |
| *A* | A3 | -0.008 | 0.988 | 122.240 | 59.8 | 2.44E-02 | 43.6 | 9.29E-03 | 267 | 1.28E-03 | 2.04 |
| *A* | A3 | -0.008 | 0.968 | 127.087 | 52.4 | 9.67E-03 | 40.8 | 4.09E-03 | 208 | 1.08E-03 | 2.43 |
| *A* | A3 | -0.011 | 0.957 | 95.200 | 47.6 | 2.67E-02 | 29.9 | 1.13E-02 | 192 | 2.96E-03 | 2.00 |
| *A* | A3 | -0.009 | 0.991 | 113.470 | 52.4 | 1.42E-02 | 39 | 5.81E-03 | 241 | 1.11E-03 | 2.17 |
| *B* | B1 | -0.005 | 0.968 | 219.243 | 62.8 | 1.85E-02 | 40.8 | 7.40E-03 | 368 | 9.13E-04 | 3.49 |
| *B* | B1 | -0.009 | 0.969 | 108.305 | 20.9 | 3.48E-03 | 52.4 | 2.91E-03 | 152 | 4.43E-04 | 5.18 |
| *B* | B1 | -0.009 | 0.969 | 108.051 | 29.8 | 3.41E-03 | 52.8 | 2.72E-03 | 151 | 4.38E-04 | 3.63 |
| *B* | B1 | -0.003 | 0.964 | 371.928 | 62.8 | 2.57E-03 | 33.6 | 2.40E-03 | 388 | 2.73E-04 | 5.92 |
| *B* | B1 | -0.004 | 0.948 | 235.246 | 57.6 | 1.35E-02 | 37.2 | 5.16E-03 | 367 | 7.83E-04 | 4.08 |
| *B* | B2 | -0.006 | 0.993 | 178.572 | 52.4 | 2.45E-03 | 40.3 | 1.16E-03 | 264 | 1.42E-04 | 3.41 |
| *B* | B2 | -0.005 | 0.987 | 219.298 | 43 | 3.68E-03 | 35.1 | 2.34E-03 | 339 | 2.16E-04 | 5.10 |
| *B* | B2 | -0.003 | 0.949 | 329.515 | 47.2 | 1.07E-02 | 59.3 | 7.68E-03 | 496 | 6.39E-04 | 6.98 |
| *B* | B2 | -0.004 | 0.984 | 245.949 | 47.6 | 8.18E-03 | 62.8 | 5.49E-03 | 419 | 4.48E-04 | 5.17 |
| *B* | B2 | -0.006 | 0.996 | 180.269 | 48.3 | 2.14E-03 | 29.2 | 7.59E-04 | 236 | 1.74E-04 | 3.73 |
| *B* | B3 | -0.009 | 0.972 | 114.821 | 39.8 | 1.40E-02 | 23.6 | 1.28E-02 | 199 | 2.40E-03 | 2.88 |
| *B* | B3 | -0.011 | 0.971 | 87.641 | 62.8 | 3.97E-03 | 32.9 | 1.92E-03 | 146 | 3.70E-04 | 1.40 |
| *B* | B3 | -0.007 | 0.999 | 151.059 | 23 | 1.19E-03 | 43.6 | 1.07E-03 | 261 | 1.91E-04 | 6.57 |
| *B* | B3 | -0.006 | 0.991 | 159.499 | 62.8 | 5.96E-02 | 43.6 | 1.02E-03 | 228 | 2.43E-04 | 2.54 |
| *B* | B3 | -0.013 | 0.917 | 75.420 | 43 | 1.70E-02 | 18.6 | 1.65E-02 | 102 | 3.29E-03 | 1.75 |
| *C* | C1 | -0.014 | 0.966 | 70.757 | 21.1 | 8.25E-03 | 31.1 | 4.09E-03 | 138 | 1.29E-03 | 3.35 |
| *C* | C1 | -0.004 | 0.991 | 244.913 | 39.8 | 1.28E-02 | 26.2 | 5.58E-03 | 468 | 7.64E-04 | 6.15 |
| *C* | C1 | -0.008 | 0.994 | 122.542 | 62.8 | 1.27E-02 | 25.9 | 7.72E-03 | 169 | 1.57E-03 | 1.95 |
| *C* | C1 | -0.004 | 0.968 | 249.356 | 47.2 | 1.09E-02 | 58.2 | 9.50E-03 | 535 | 6.75E-04 | 5.28 |
| *C* | C1 | -0.005 | 0.968 | 203.062 | 49.5 | 3.07E-02 | 31.3 | 1.79E-02 | 499 | 1.86E-03 | 4.10 |
| *C* | C2 | -0.012 | 0.728 | 85.180 | 55.6 | 1.15E-02 | 34 | 5.02E-03 | 356 | 8.19E-04 | 1.53 |
| *C* | C2 | -0.003 | 0.960 | 309.358 | 27.2 | 2.57E-02 | 25 | 1.61E-02 | 833 | 1.88E-03 | 11.37 |
| *C* | C2 | -0.005 | 0.954 | 189.840 | 54.2 | 6.05E-02 | 41.3 | 2.16E-02 | 629 | 2.02E-03 | 3.50 |
| *C* | C2 | -0.003 | 0.943 | 300.523 | 59.8 | 7.55E-03 | 44.9 | 7.53E-03 | 594 | 6.88E-04 | 5.03 |
| *C* | C2 | -0.004 | 0.938 | 243.539 | 40.3 | 3.57E-03 | 27.7 | 1.77E-03 | 716 | 2.77E-04 | 6.04 |
| *C* | C3 | -0.006 | 0.985 | 166.940 | 50.3 | 1.53E-02 | 28.3 | 4.77E-03 | 291 | 9.91E-04 | 3.32 |
| *C* | C3 | -0.007 | 0.981 | 147.955 | 54.2 | 1.50E-02 | 34.1 | 1.38E-02 | 283 | 1.53E-03 | 2.73 |
| *C* | C3 | -0.006 | 0.978 | 154.572 | 28.3 | 8.72E-03 | 53.7 | 6.69E-03 | 332 | 8.96E-04 | 5.46 |
| *C* | C3 | -0.006 | 0.978 | 171.765 | 48 | 1.34E-02 | 29.6 | 1.28E-02 | 329 | 1.64E-03 | 3.58 |
| *C* | C3 | -0.006 | 0.979 | 154.984 | 25.5 | 4.85E-03 | 30.8 | 4.64E-03 | 362 | 6.67E-04 | 6.08 |

Footnote: The full column names from left are colony, polyp, exponent from model fit, R-squared value, time constant of mixing (sec), primary peak period (sec), primary peak power ($ppm^2$), secondary peak period (sec), N (sampling rate of 1 Hz), integrated power of the power spectral density ($ppm^2$ radians$^{-1}$), and ration of tau to the primary period.
